# A transphyletic study of metazoan β-catenin protein complexes

**DOI:** 10.1101/2024.07.01.600363

**Authors:** Ivan Mbogo, Chihiro Kawano, Ryotaro Nakamura, Yuko Tsuchiya, Alejandro Villar-Briones, Yoshitoshi Hirao, Yuuri Yasuoka, Eisuke Hayakawa, Kentaro Tomii, Hiroshi Watanabe

**Affiliations:** Evolutionary Neurobiology Unit, Okinawa Institute of Science and Technology Graduate University, Okinawa, Japan; Artificial Intelligence Research Center, National Institute of Advanced Industrial Science and Technology (AIST), Tokyo, Japan; Instrumental Analysis Section, Okinawa Institute of Science and Technology Graduate University, Okinawa, Japan; Marine Genomics Unit, Okinawa Institute of Science and Technology Graduate University, Okinawa, Japan

**Author notes:** Ivan Mbogo, Sysmex Corporation, Ltd. 1-5-1, Chuo-ku, Kobe, 651-0073, Japan. Eisuke Hayakawa, Department of Bioscience and Bioinformatics, Kyushu Institute of Technology, 680-4, Kawazu, Iizuka, Fukuoka, 820-8502, JAPAN. Alejandro Villar-Briones, Project Planning and Implementation Section, Okinawa Institute of Science and Technology Graduate University, Okinawa, Japan. Yuuri Yasuoka, Laboratory for Comprehensive Genomic Analysis, RIKEN Center for Integrative Medical Sciences, Yokohama, Japan. Corresponding author: Correspondence to Hiroshi Watanabe.

**Keywords:** β-catenin, metazoan, Nematostella vectensis, protein complex

## Abstract

β-catenin is essential for various biological processes, such as body axis determination and cell differentiation, during embryonic development in metazoans. β-catenin functions are thought to be exerted through complexes formed with various types of proteins. Although β-catenin complex proteins have been identified in several bilaterians, little is known about the structural and functional properties of β-catenin complexes in early metazoan evolution. In this study, we performed a comparative analysis of β-catenin sequences in nonbilaterian lineages that diverged early in metazoan evolution. We also carried out transphyletic function experiments with β-catenin from nonbilaterian metazoans using developing *Xenopus* embryos, which included secondary axis induction in embryos and proteomic analysis of β-catenin protein complexes. Comparative functional analysis of nonbilaterian β-catenins demonstrated sequence characteristics important for β-catenin functions, and the deep origin and evolutionary conservation of the cadherin-catenin complex. Proteins co-immunoprecipitated with β-catenin included several proteins conserved among metazoans. These data provide new insights into the conserved repertoire of β-catenin complexes.

## Background

Recent genome analysis of early-branching, nonbilaterian metazoans, including cnidarians, poriferans, and ctenophores, showed that many genes involved in β-catenin signaling were acquired in parallel with the evolutionary emergence of the Metazoa. In Cnidaria, the closest sister lineage to Bilateria, the oral-aboral axis is the main body axis, and considerable evidence from research on selected cnidarians indicates that canonical Wnt signaling involving the β-catenin/TCF pathway exhibits high activity on the oral side and controls oral-aboral axis formation [1–5] Indeed, in *Nematostella vectensis* (Anthozoa, Cnidaria), transplantation of a fragment from the blastopore lip, which has high β-catenin activity and will later become the mouth, to the aboral ectoderm, induces a secondary body axis [6]. Similarly, transplantation of the hypostome to a different site causes a secondary body axis in *Hydra vulgaris* (Hydrozoa, Cnidaria) [7]. These organizer functions of the blastopore/hypostome can be mimicked by enhancing β-catenin signals by treatment with a GSK3β inhibitor [8,9]. In addition, β-catenin binds to and co-localizes with the cadherin-catenin cell adhesion complex in *Nematostella* [10,11], as it does in bilaterians. In ctenophores and poriferans, gene components of β-catenin signaling are localized at particular positions along the developing embryo axis [12–15].

Data showing that multiple functions of β-catenin are already functional in early-branching metazoan lineages suggest that β-catenin’s diverse protein complex repertoire was established, at least in part, early during metazoan evolution. Actually, a pioneering study in *Ephydatia muelleri* (Porifera), showed that cadherins and α-catenin bind to endogenous β-catenin, suggesting that the core complex of β-catenin machinery involved in cell-cell adhesion has deep evolutionary roots [16]. However, neither the complete picture of protein complexes formed by early β-catenin species nor commonalities in their components are known. To date, the β-catenin complex repertoire has been analyzed mainly in mammalian cell lines/tissues, including studies employing immunoprecipitation with mass spectrometry (IP-MS) [17, 18]. Recently, in studies of β-catenin protein-protein interactions (PPIs), researchers have begun to move away from cell cultures to cultured tissues, allowing identification of tissue-specific PPIs [19,20].

To compare proteins that bind to β-catenin in various early-branching metazoans, we employed a transphyletic experimental approach, in which we examined proteins of the complex formed by heterologous β-catenins expressed in developing *Xenopus* embryos. The transphyletic approach is classical, but still constitutes an effective technique for testing evolutionary conservation of protein functions and gene-regulatory networks, especially in comparative analyses of diverse phyla, including non-model early-branching nonbilaterians [21–26]. This system also allows an experimental comparison among β-catenins from phylogenetically distant species for β-catenin complexes associated with specific biological contexts.

In this study, to gain insights into evolution of the β-catenin protein complex, we first performed a comparative analysis of β-catenin sequences from nonbilaterian lineages that diverged early in metazoan evolution. Especially amino acid residues and motifs related to β-catenin function in bilaterians were examined in nonbilaterian β-catenins. Next, we performed a transphyletic experiment with β-catenin derived from a non-bilaterian metazoan using developing *Xenopus* embryos, one of the best experimental systems for verifying organizer-inducing activity of β-catenin. Finally, we carried out a proteomic survey of proteins immunoprecipitated with nonbilaterian β-catenin from *Xenopus* gastrula embryos to provide a potential list of evolutionarily conserved protein components of the β-catenin machinery.

## Results

### Sequence comparisons and 3D structures of metazoan β-catenins

In analyzing early evolution of the β-catenin complex, we first examined conservation of β-catenin sequences of early-branching nonbilaterians (Fig. 1). This is important for understanding post-translational modifications and direct protein interaction sites. As a reference structure, we included ARM6, a β-catenin-like protein from the choanoflagellate, *Salpingoeca rosetta*. Choanoflagellates, are the sister group of Metazoa. Choanoflagellate ARM6 is ancestral to metazoan β-catenin and Adenomatous Polyposis Coli (APC) groups and has fewer armadillo repeats (Fig. 1A, B) [27]. Overall, cnidarian and poriferan β-catenins share many functional motifs with bilaterian β-catenin, but the proportion decreases in ctenophores, and except for a few phosphorylation sites, most functional motifs are not conserved in ARM6 of Choanoflagellate.

**Figure 1.**
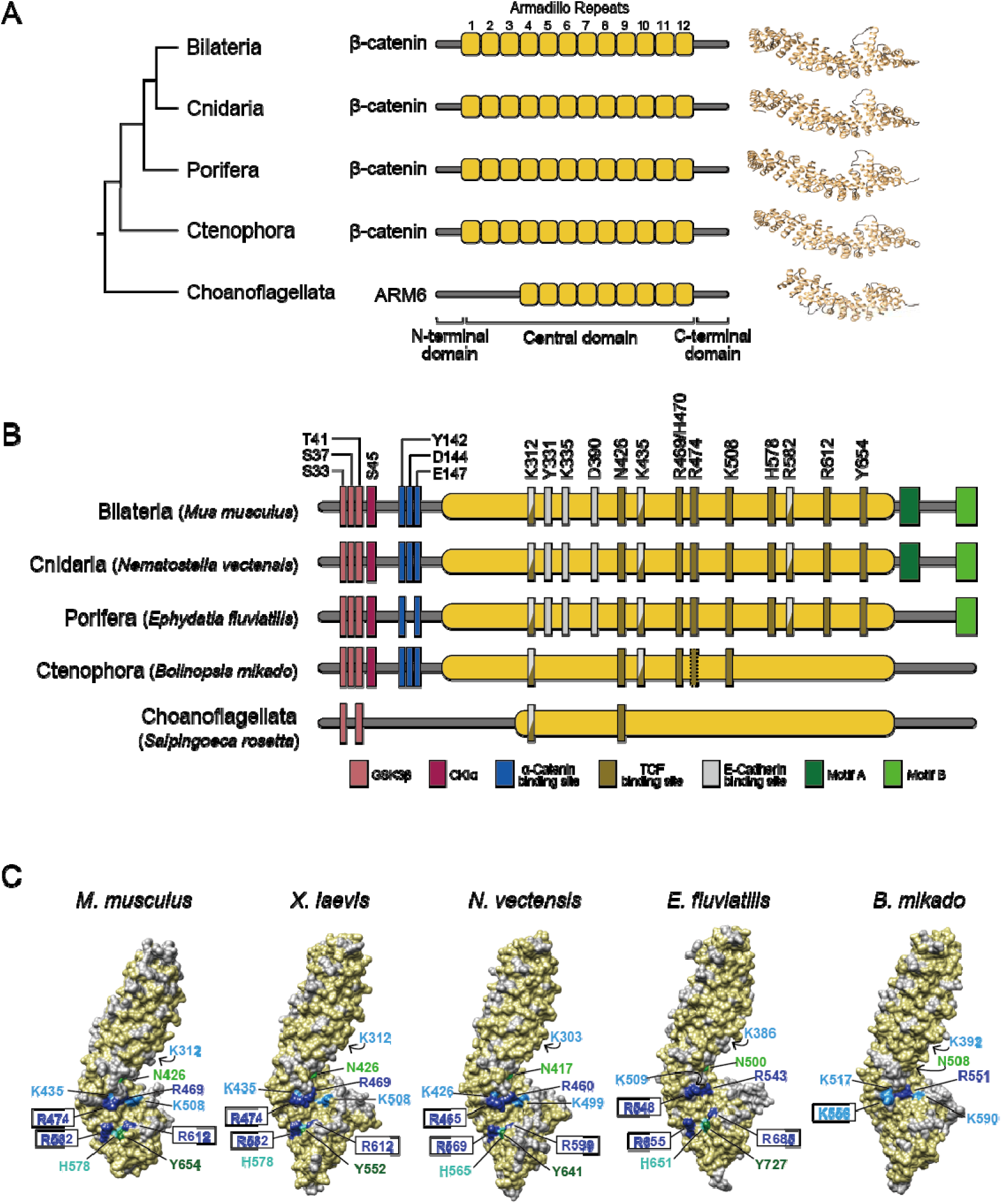
Primary and 3D structures of nonbilaterian β-catenins. (A) Overview of phylogeny and structure of β-catenins of early-branching (nonbilaterian) metazoans and ARM6 protein of the unicellular sister, Choanoflagellata. B) Domain organizations of β-catenin proteins. Metazoan β-catenins show a high degree of conservation of the central region, GSK3β/CK1α phosphorylation sites, and αE-catenin binding sites. At the C-terminus, motif A is conserved in only Bilateria and Cnidaria, whereas motif B is shared among all metazoan lineages, except for Ctenophora. In Choanoflagellata, *Salpingoeca rosetta*, possible GSK3β phosphorylation sites (S33/T41) exist. (B) Surface depictions of β-catenin homology structural models were compared with mouse β-catenin. This confirmed that the three crucial arginine residues (dark blue) of nonbilaterians were localized similarly to mouse and *Xenopus* β-catenins. K556, R551, and K590 of *B. mikado* β-catenin were located at positions corresponding to mouse β-catenin R474, R469, and K508, but the R582, H578, R612, Y654 assembly of mouse β-catenin is completely absent in *B. mikado* β-catenin. Structure visualized using UCSF Chimera software.

In our comparison, GSK3β and CK1α phosphorylation sites critical for β-catenin degradation are apparently conserved in all metazoans, except for *Hydra*, which lacks the site corresponding to S37 (Fig. 1B, S1, S2). Interestingly, in unicellular *S. rosetta*, two residues corresponding to S33 & T41 were also conserved. The central domain is the leading interaction site for several proteins, including TCF and Cadherin1(CDH1) [28, 29]. A high degree of conservation was observed in Bilateria, Cnidaria, and Porifera in the central region (Fig. 1B). Cnidarian and poriferan β-catenins displayed full conservation of the three critical lysine residues (K312, K335, and K435 in mouse β-catenin) and others (Y331, D390, and R582) that are required for E-cadherin binding [28, 29]. Among these sites, only two lysines, K312 and K435, were conserved in Ctenophora (Fig. 1B, S1).

**Figure 2.**
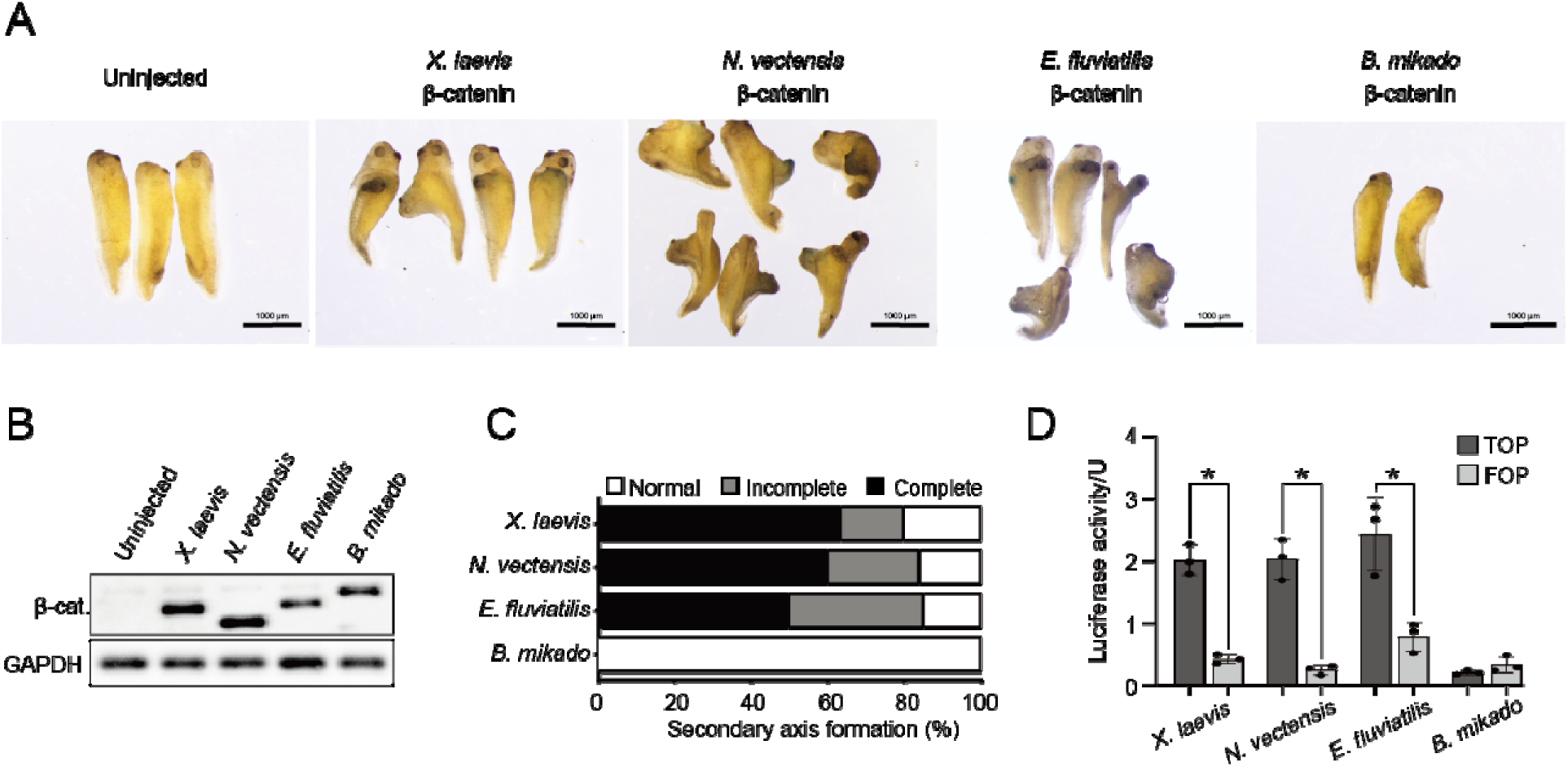
*Xenopus* secondary axis induction by nonbilaterian β-catenins. (A) Cnidarian (*N. vectensis*) and poriferan (*E. fluviatilis*) β-catenins (100 pg mRNA) induced a secondary body axis similar to that induced by *Xenopus* β-catenin. Ctenophore (*B. mikado*) β-catenin had no inductive activity. (B) Western blot analysis of Flag-tagged β-catenins confirmed the expression of nonbilaterian β-catenin proteins in *Xenopus* embryos. (C) The majority of secondary body axes induced by injection of *X. laevis* (n = 24), *N. vectensis* (n = 20), and *E. fluviatilis* (n = 18) β-catenin mRNAs (100 pg) were complete (the second axis had a pair of eyes and a cement gland). This was followed by incomplete axis inductions in which head features were not fully developed. Secondary axes were not observed in embryos injected with *B. mikado* β-catenin mRNA (100 pg) (n = 19). (D) Expression of *X. laevis, N. vectensis*, and *E. fluviatilis* FLAG-tagged β-catenin mRNA (100 pg) in *Xenopus* embryos resulted in a significant increase in β-catenin TOPflash activity. No detectable level of TOPflash activation was observed by expression of *B. mikado* β-catenin. Asterisks denote statistical significance *p* <0.0001 (Two-way ANOVA). The result is representative of two reproducible, independent experiments.

Three arginine residues of mouse β-catenin, R474, R582, and R612, are important in binding to TCF [28, 30]. These amino acids are conserved in Cnidaria and Porifera, but in Ctenophora, the residue corresponding to R474 is replaced with a lysine residue, and R582 & R612 are not conserved. To gain further insight into the influence of these differences on binding of TCF, we investigated other amino acids, K312, N426, K435, R469, H470, K508, H578 and Y654, that are thought to be involved in β-catenin-TCF interactions in mice [28, 30, 31] (Fig. 1B, S1). Like the three important arginine residues (R474/R582/R612) mentioned above, these amino acids are also common among Bilateria, Cnidaria, and Porifera, and all except H578 and Y654 are conserved in Ctenophora. Interestingly, most of these amino acids that function in binding of β-catenin to TCF are not conserved in *S. rosetta*. However, *S. rosetta* ARM6 has asparagine corresponding to N426 and arginine at a position corresponding to lysine (K312), which has chemical properties similar to those of lysine. The C-terminus of β-catenin, which has a transactivation domain, is important for regulation of gene expression by the β-catenin/TCF complex [32, 33]. Its signaling ability is probably due to two motifs, A and B [33–36]. Alignment combined with Multiple Expectation maximization for Motif Elicitation (MEME) analysis confirmed that motif A is present only in Bilateria and Cnidaria, whereas motif B is also present in Porifera (Fig. 1B, S1, S3). On the other hand, neither motif A or B exists in ctenophore β-catenin and choanoflagellate ARM6.

**Figure 3.**
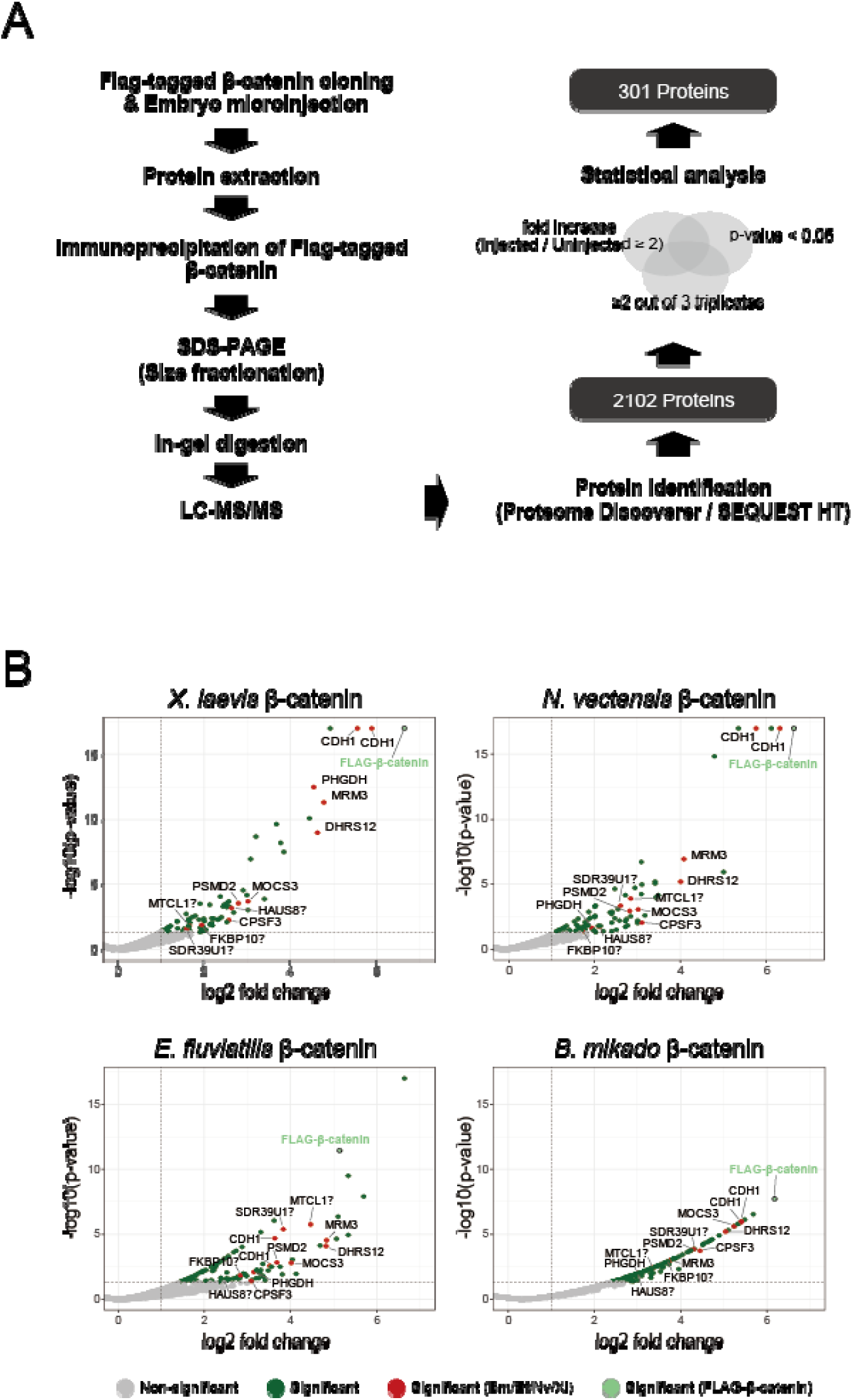
Precipitation of β-catenin interacting proteins. (A) Scheme of our IP-MS analysis of β-catenin protein complexes. (B) Mass spectrometry volcano plots resulting from analysis of enriched metazoan FLAG-tagged β-catenins expressed in *Xenopus* embryos. High confidence proteins (1% FDR), with a fold abundance ratio ≥2 and a p-value <0.05, were considered “true” interactions.

Finally, we compared amino acids required for binding to α-catenin. The α-catenin binding site in β-catenin was previously narrowed down to amino acids 118-146 (in mice) [37].

Metazoan-wide conservation across this region is not very high, except for a few residues. In mouse β-catenin, Y142 is vital for α-catenin binding, since mutation to alanine eliminates β-catenin-α-catenin interaction [37]. The site corresponding to Y142 is conserved in all metazoans (Fig. 1B, S1, S4). Two acidic residues, D144 and E147, also affect interaction with α-catenin [38]. D144 is not conserved among poriferans. *S. rosetta* has a large insertion sequence in this region, which precluded accurate verification.

**Figure 4.**
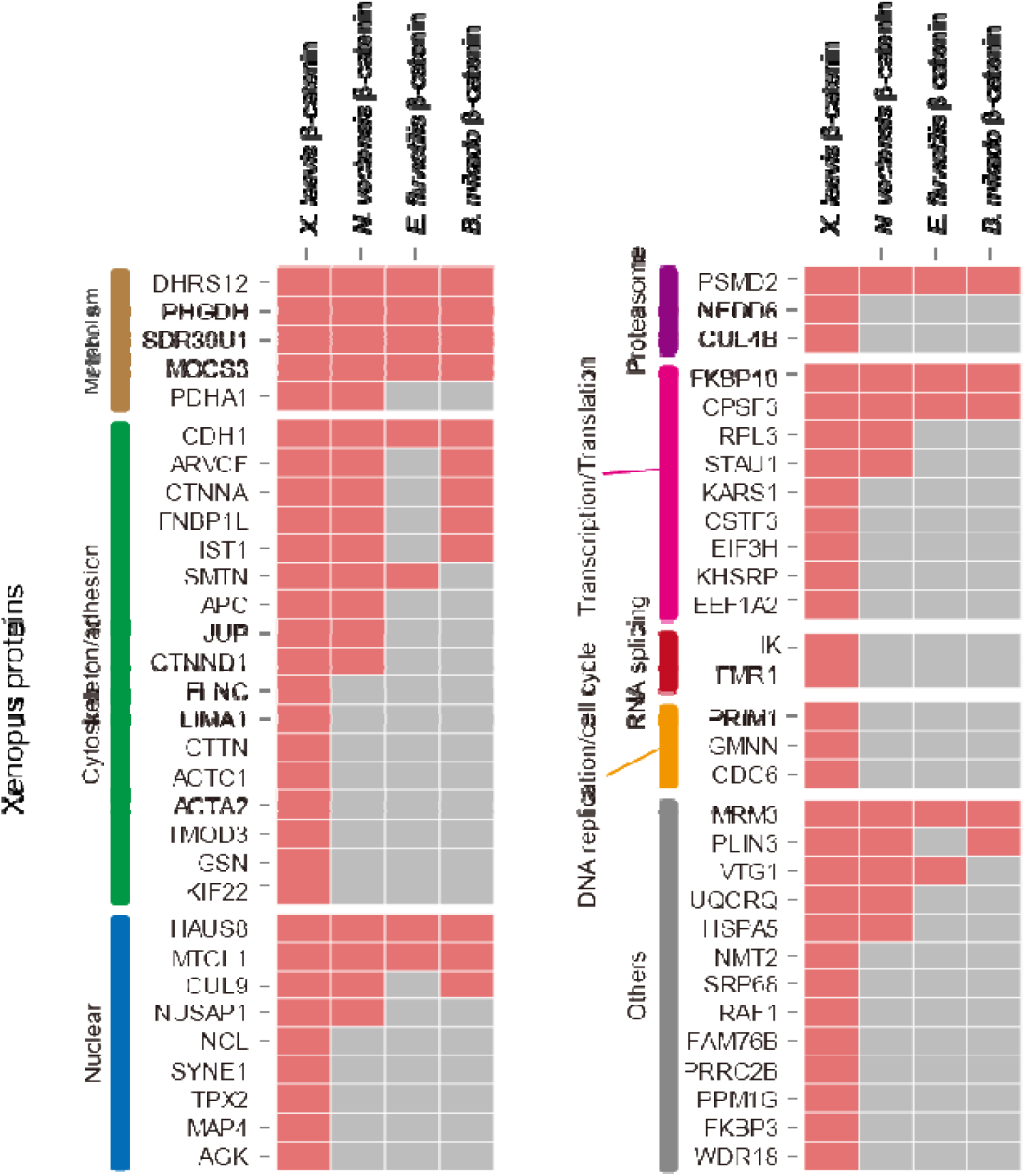
Potential evolutionarily conserved β-catenin interactions. Several proteins, e.g., CDH1, ARVCF, CTNNA1, that interact with β-catenin mainly in bilaterian models, formed complexes with basal metazoan β-catenins. Furthermore, new interactions were identified that may shed additional light on functional evolution of β-catenin protein machinery in metazoans.

β-catenin sequences from *Xenopus laevis* (Vertebrata), *Nematostella vectensis* (Cnidaria), *Ephydatia fluviatilis* (Porifera)*, Bolinopsis mikado* (Ctenophora), and ARM6 of *S. rosetta* were then used in homology modeling with the crystal structure of mouse β-catenin (PDB ID: 4ev8) to investigate structural evolution. Phi & Psi distributions of Ramachandran plots generated with PROCHECK indicated that over 90% of amino acids in modeled structures were in favored positions (Fig. S5). Despite amino acid variations of β-catenin observed among metazoans (Fig. S6), the structure of the predicted armadillo repeat region of β-catenin appears to be stable since emergence of the Metazoa. Interestingly, between residues 452-878, the *S. rosetta* ARM6 protein possesses a structure that resembles metazoan β-catenins (Fig. S7). However, *S. rosetta* ARM6 has a short loop, whereas metazoan β-catenins have a long loop between armadillo repeats 10 and 11, indicating that this is probably a metazoan invention (Fig. 1A). The armadillo repeat region contains multiple amino acids that are or may be involved in TCF binding regions (Fig. 1B). These amino acids are conserved in β-catenins from nonbilaterians. We also demonstrated steric conservation of these amino acids. Thus, in addition to sequence similarities, orientations of TCF-binding amino acids are highly conserved in β-catenin from Bilateria, Cnidaria, and Porifera (Fig. 1C). In Ctenophora, homology was observed only in a few amino acids.

Given their structural similarities, the next question is, “To what extent are functions of β-catenins conserved?” Functional analysis of nonbilaterian β-catenins, using mainly *Nematostella*, shows that β-catenin is important in the blastoporal organizer and during gastrulation and subsequent endoderm fate determination. On the other hand, in poriferans and ctenophores, knowledge of β-catenin function is much more limited [12, 15, 16, 39]. During developmental of *Mnemiopsis leidyi* (Ctenophora), perturbation of classical β-catenin signaling had limited effect [12]. Classical β-catenin complexes have also been reported in adult *Ephydatia*, but their functions remain unknown [16].

### Comparison of axis-inducing activity of metazoan **β**-catenins in *Xenopus* embryos

To investigate conservation of β-catenin activity, we performed a secondary axis induction assay using an ectopic expression system with *Xenopus* embryos. *In vivo* embryonic systems, which require a variety of cellular developmental events such as epithelial formation, active cell division, and formation of signal centers necessary for body axis patterning, are ideal for comprehensive functional surveys of the β-catenin complex.

Differences in codons in mRNAs of different organisms can affect rates of translation when expressed in other organisms. Therefore, we began by optimizing the amount of mRNA injected to achieve expression of relatively similar levels of FLAG-tagged proteins (Fig. S8A). For all metazoan β-catenins, FLAG-tagged β-catenin protein was observed at the expected band sizes. However, expression of *S. rosetta* ARM6 in *Xenopus* could not be confirmed even when the amount of injected mRNA was increased (Fig. S8B). Therefore, the following analysis was performed using only metazoan β-catenins.

To test whether basal metazoan β-catenins are functional, they were injected into the ventral equatorial region of one blastomere of *Xenopus* 4-cell embryos. Figure 2B shows that injection of 100 pg mRNA resulted in expression of FLAG-tagged β-catenin proteins. These results showed that *N. vectensis* and *E. fluviatilis* β-catenin induced a secondary body axis similar to that induced by *X. laevis* β-catenin (Fig. 2A-C). In contrast, *B. mikado* β-catenin did not induce a secondary axis. Even when *B. mikado* β-catenin mRNA was injected at high concentrations (>500 pg), only protrusion-like structures without head characteristics appeared in some embryos (Fig. S9). This suggests that the lack of secondary axis-inducing ability of *B. mikado* β-catenin is due to differences in signals it can activate, rather than to weakness of activation.

To further confirm whether basal metazoan β-catenins can drive β-catenin/TCF signaling, a TOPflash luciferase assay using *Xenopus* embryos was performed [40, 41]. Consistent with secondary axis induction experiments, there was a significant increase in luciferase activity following expression of *X. laevis*, *N. vectensis*, and *E. fluviatilis* β-catenin (Fig. 2D), confirming that signaling activity of β-catenin is conserved in Cnidaria and Porifera. On the other hand, expression of *B. mikado* β-catenin did not increase luciferase activity. This reflects reduced conservation of amino acids necessary for TCF binding in ctenophore β-catenins (Fig. 1B). Additionally, together with the finding that β-catenin inhibition in *Mnemiopsis* does not clearly affect conserved functions, such as body axis formation [12], these data suggest that ctenophore β-catenin does not contribute significantly to canonical Wnt signaling (Wnt/β-catenin/TCF signaling).

The capacity of cnidarian and poriferan β-catenins to induce phenotypes similar to that of *Xenopus* β-catenin suggests that they all form a common protein complex in *Xenopus* embryos. Therefore, we next examined the protein complex formed by each β-catenin in developing *Xenopus* embryos.

### Proteomic analysis of **β**-catenin protein complexes

To investigate protein complexes made by each basal metazoan β-catenin in developing *Xenopus* embryos, FLAG-tagged β-catenin was immunoprecipitated from homogenates at gastrula stage 11, and proteomic analysis of resulting protein complexes was performed. Western blotting confirmed that comparable amounts of exogenous FLAG-tagged β-catenin proteins were immunoprecipitated (Fig. S10). Subsequently, interacting proteins were identified using liquid chromatography with tandem mass spectrometry. An IgG control analysis from uninjected embryos was also included. To reduce false positives in identified proteins, only proteins with an abundance ratio greater than or equal to 2 (p-value < 0.05) are generally used as a threshold for “true” interactions. The presence of known bilaterian β-catenin interacting proteins confirmed that identified proteins represented “true” interactions (Fig. 3A, Table S1). Figure 3B shows a schematic of proteins co-immunoprecipitated with each β-catenin species. A number of proteins, including Cadherin (CDH), have been identified as interacting partners with all exogenous metazoan β-catenins. Our BLAST search confirmed that many of them have protein homologs in nonbilaterian metazoans.

Unexpectedly, many metabolism-related proteins, such as DHRS12, PHGDH, SDR39U1, and MOCS3, were commonly detected in bilaterian/nonbilaterian β-catenin complexes (Fig. 4). Although there are no reports that these enzymes interact with β-catenin, and their physiological significance in the β-catenin complex is currently unknown, overexpression of DHRS12 inhibits β-catenin signaling in human cell lines [42]. Another group of highly conserved components of the β-catenin complex includes proteins associated with cell adhesion. Since conservation of CDH1 binding sites on β-catenin is very high among metazoans (Fig. 1B, Fig. S1), it is not surprising that CDH1 was precipitated by all β-catenins.

Interestingly*, E. fluviatilis* β-catenin was unable to immunoprecipitate *X. laevis* α-catenin at detectable levels. This was also confirmed in western blotting analysis (Fig. S11). Poriferan β-catenins do not have the conserved amino acids required for binding to α-catenin (Fig. S4). On the other hand, binding of endogenous β-catenin and α-catenin has been observed in the poriferan, *Ephydatia muelleri* [16]. This suggests that poriferan ancestors evolved a unique interaction of β– and α-catenin.Given that β-catenin proteins in a broad array of nonbilaterians bind α-catenin and phylogenetically distant CDH1 proteins, indicates that β-catenin complex functions involved in the adherens junction were acquired among the earliest metazoans (even if there were unique modifications in the Porifera) and remain highly conserved. Since organizer-inducing activity was observed in bilaterian, cnidarian, and poriferan β-catenins in developing *Xenopus* embryos, we expected to detect protein complexes specific to these β-catenin immunoprecipitates. However, proteomic analysis detected only a few proteins common to bilaterian/cnidarian/poriferan β-catenins. Smoothelin (SMTN) was identified as a binding protein common to β-catenin of *X. laevis*, *N. vectensis*, and *E. fluviatilis*. In bilaterians, SMTN binds Cortactin (CTTN) and stabilizes the cortical actin meshwork of epithelial cell membrane [43]. Cortactin binds β-catenin, α-catenin, and p120 catenin (catenin delta-1/ CTNND1) and is important for the function of adherens junctions [44]. VTG1, which has an unknown function, was also detected in this group; however, even though gene homologs of SMTN and VTG1 exist in *X. laevis* and *N. vectensis*, they are not in the genome of *E. fluviatilis* (Fig. S12).

## Discussion

Several studies have suggested that β-catenin was involved in the rise of multicellularity and further evolution of complex body organization. However, little is known about structural attributes of β-catenin that underlie its various functions, and the evolutionary process of β-catenin machinery. In this study, in addition to detailed sequence analysis of metazoan β-catenins, we utilized structural, proteomic, and functional analyses to understand evolutionary dynamics of the β-catenin complex.

### Conserved and lineage-specific features of **β**-catenin cell adhesion complexes

Key lysine residues (K312 and K435) for binding E-cadherin are conserved in all metazoan β-catenins, and in fact, it was confirmed that E-cadherin is co-immunoprecipitated with all metazoan β-catenins from *Xenopus* embryos. It has been suggested that Y331, K335, D390, and R582 residues of mouse β-catenin are important for binding to an intercellular domain of E-cadherin [29]. However, these residues are conserved in Cnidaria and Porifera, but not in Ctenophora. Considering that *Xenopus* E-cadherin can bind all metazoan β-catenins, including those from Ctenophora, these amino acids appear non-essential for binding to E-cadherin. The inability of poriferan β-catenin to bind *Xenopus* α-catenin was surprising, as previous research had confirmed endogenous Interaction of these proteins in *Ephydatia* [16]. The cause appears to be the charge distribution near the critical tyrosine residue in the α-catenin binding site. In mouse β-catenin, the hydroxyl group of Y142 is close to two acidic amino acids, D144 and E147. Mutating these residues or introducing a negative charge on Y142 could alter charge distribution, preventing α-catenin binding [38]. In both *Ephydatia muelleri* and *Ephydatia fluviatilis*, acidic residues are conserved at the site corresponding to E147, whereas D144 is substituted for the neutral glycine, possibly impeding, or weakening α-catenin binding. This implies that poriferan β-catenin and α-catenin co-evolved to either allow or strengthen interaction. Details of molecular features of the cell adhesion system of poriferan epithelia have not yet been clarified. Furthermore, in Ctenophora, genetic peculiarities of elements involved in cell adhesion machinery have been noted [29]. Understanding functions of cadherins and α-catenins in Ctenophora and Porifera remains for further study.

### Organizer induction and **β**-catenin signaling

The key to explaining the organizer induction results might have been co-immunoprecipitation of TCF with β-catenin, but our IP-MS did not identify any TCF peptides. This is probably because the β-catenin/TCF complex is not very stable. Indeed, previous β-catenin IP-MS studies have also been unsuccessful at detecting β-catenin-TCF interaction [18, 20, 45]. Further experimental optimization is needed to discover the molecular organization of the transcription regulatory machinery formed by β-catenin and transcription factors such as TCF, but comparisons of the function and sequence of β-catenin in basal metazoans still provided new insights. Some amino acids of β-catenin that are key to binding TCF are conserved in all metazoans, but four sites corresponding to R582, H578, R612, and Y654 in mouse β-catenin are not conserved in ctenophores. However, a previous study using *X. laevis* β-catenin showed that mutations R612A & Y654A do not significantly affect TCF interaction [31]. These may, therefore, not be evolutionarily critical in binding TCF. We also observed domain differences in the C-termini of ctenophore β-catenins. Previous studies have shown that β-catenin C-termini can function as transcriptional activators when fused to TCF [46, 47]. Furthermore, a β-catenin C-terminus-LEF1 fusion is sufficient for secondary body axis induction in *Xenopus* [32]. Two motifs, A and B, have previously been identified in the C-terminus [34]. Motif A was confined to bilaterians and cnidarians. It is localized in the Helix-C region, a key region for β-catenin transcription activity [48]. However, motif A is probably not essential for organizer induction, as it is absent in *E. fluviatilis* β-catenin that still managed to drive secondary axis induction. Our focus then turned to motif B, which is present in all metazoans except ctenophores. A recent study showed that mutation of this domain resulted in decreased TCF-dependent transcriptional activity [35]. Therefore, organizer induction capacity may depend on the interaction at R582/H578 and motif B with transcription factors such as TCF. The inability of *B. mikado* β-catenin to induce a secondary body axis could be because it cannot fully activate downstream genes of β-catenin/TCF, possibly due to insufficient interaction at motif B with endogenous proteins.

### **β**-catenin and microtubule functions

Our study uncovered several centrosome-related proteins. β-catenin centrosomal localization has been observed in *Caenorhabditis elegans* (Nematoda) and *Platynereis dumerilii* (Annelida) [49–52]. However, although it has been suggested that centrosomal accumulation is conserved, there is no evidence for this in nonbilaterians.

HAUS8, now identified as a member of the β-catenin complex, is part of a multi-subunit protein complex that controls centrosome and spindle integrity [53]. Depletion of HAUS8 resulted in centrosome defects, delayed mitosis, and increased aneuploidy [54]. Since β-catenin localization at centrosomes is associated with mitotic progression [55], it is plausible that HAUS8 and β-catenin interact to promote mitosis. The HAUS8 gene is absent before the Bilateria and Cnidaria, and it may be a new interaction adopted by these common ancestors to control cell division.

Several other β-catenin partners are common to Bilateria and Cnidaria. Of interest among these is a group of proteins essential for microtubule formation. One of them, APC, forms a complex with β-catenin in bilaterians [56, 57]. As expected, we found that *Xenopus* APC binds to the complex of *X. laevis* and *N. vectensis* β-catenin proteins. APC was previously detected at centrosomes in interphase and mitosis [58, 59]. Recent expression studies showed that APC functions at centrosomes by stimulating microtubule growth, since expression of a truncated APC reduced the rate of microtubule growth [60]. Finally, a microtubule-associated protein, NuSAP1, was identified as interacting with both *X. laevis* and *N. vectensis* β-catenin. NuSAP1 activates β-catenin signaling and its depletion results in abnormal mitotic spindles, followed by abnormal chromosome segregation and cytokinesis [61, 62]. These proteins may have been critical in strengthening β-catenin functionality in the last common ancestor of the Bilateria and Cnidaria.

## Conclusions

In this study, we conducted a comprehensive analysis of sequences, structures, and binding protein repertoires of β-catenins, a metazoan invention that is responsible for the oldest multifunctional signal pathway. We particularly focused on comparisons using primary and 3D structure predictions, and transphyletic functional analysis, using developing *Xenopus* embryos, a biological context in which β-catenin serves an important function. Our results show high conservation of β-catenin sequences in Bilateria/Cnidaria/Porifera, and clarify the uniqueness of β-catenin in Ctenophora, the earliest branching metazoan lineage. Although proteomic analysis did not reveal proteins associated with the functional gap between Bilateria/Cnidaria/Porifera and Ctenophora β-catenin complexes, structural and functional comparisons revealed the most evolutionarily conserved β-catenin region essential for interacting with TCF. Comparable proteomic analysis of various metazoan β-catenin complexes identified many complex constituent proteins, including novel members, and revealed their repertoire and phylogenetic distribution. These phylum-wide β-catenin complex lists provide foundational insights for future analysis to follow the evolutionary process of β-catenin function.

### Materials and Methods Structural evolution of **β**-catenin

Bidirectional protein BLAST searches using mouse β-catenin were carried out against various databases to extract metazoan β-catenin sequences from representative bilaterians (*Xenopus laevis*), Cnidaria (*Nematostella vectensis*), Ctenophora (*Bolinopsis mikado*), and Porifera (*Ephydatia fluviatilis*) and as an outgroup, *Choanoflagellata* (*Salpingoecca rosetta*). Amino acid sequences of *E. fluviatilis* and *B. mikado* were aligned with MIQS, a scoring matrix optimized to detect distant homologs [63]. The putative β-catenin of *Salpingoeca* was aligned with structurally known β-catenin using HHpred [64]. Due to a high substitution rate at the N– and C-termini, alignment was focused on the region from the α-catenin binding site to the 12^th^ armadillo repeat. Multiple sequence alignments of β-catenin were calculated using FAMSA [65]. β-catenin 3D models were built using MODELLER v9.20 [66], based on alignments of targeted, structurally known β-catenins, calculated with FORTE/ DELTA-FORTE, which are profile-profile alignment methods [67]. Structural models predicted with PROCHECK [68], which generates Ramachandran plots, showed the distribution of combinations of backbone dihedral angles Phi & Psi [69]. Visualization of final model structures was carried out using UCSF Chimera (Version 1.15) [70].

For domain analysis, all sequences were aligned using MUSCLE [71]. Manual curation was then undertaken to match the current alignment with the alignment used for structural analysis, resulting in a final high-quality alignment. To identify possible motifs at the C-terminus, MEME analysis was performed on unaligned sequences using MEME Suite 5.3.4 [72] with a maximum motif width of 20.

### Animal culture

Adult male and female *Xenopus laevis* were purchased from *Xenopus* Aquaculture for Teaching Materials (Ibaraki Prefecture, Japan) and maintained in our frog facility. All experiments with *X. laevis* were approved by the Animal Care and Use Committees at Okinawa Institute of Science and Technology Graduate University.

### Gene cloning and mRNA preparation

Coding sequences of *X. laevis*, *N. vectensis*, *E. fluviatilis*, and *B. mikado* β-catenin genes with an N-terminal FLAG-tag were cloned into pCS2+ plasmids (Table S3, S4). Following plasmid linearization, 1 µg of linearized DNA was used for mRNA synthesis with an mMESSAGE mMACHINE SP6 Transcription Kit (Ambion) following the manufacturer’s guidelines, with a few changes. Briefly, components of the transcription kit were thawed at room temperature and then kept on ice apart from the 10x Reaction buffer. Next, the reaction mix was set up by combining 10 µL 2X NTP/CAP, 2 µL 10x Reaction buffer, 1 µg linear template DNA, 2 µL enzyme mix, and an appropriate amount of nuclease-free water to a final volume of 20 µL. The reaction was mixed by pipetting and incubated for 4 h at 37°C. 1 µL TURBO DNase was then added to the reaction mix and incubated at 37°C for 20 min. Finally, a MEGAclear Transcription Clean-Up Kit (Ambion) was used to purify synthesized mRNA following the manufacturer’s instructions. The mRNA concentration was measured using a Nanodrop 2000c spectrophotometer (Thermo Fisher Scientific, Inc), and RNA quality was confirmed using an Agilent 4200 Tapestation (Agilent). The mRNA was then aliquoted and stored at –80°C.

### Secondary axis induction assay

*X. laevis* embryos were obtained by *in vitro* fertilization [73] and staged according to Nieuwkoop and Faber [74]. mRNA (100 pg) was co-injected with tracer LacZ mRNA (encoding β-galactosidase) (25 pg) into the ventral equatorial region of one blastomere at the 4-cell stage. Injected embryos were then incubated overnight at 20°C in 1x Steinberg solution containing 5% Ficoll and then transferred into 0.1x Steinberg until stage 37, followed by X-gal staining. Axis duplication success was scored as “Complete” if the second axis had eyes and a cement gland, “Incomplete” if head features of the secondary axis were not fully developed, and “Normal” if there was only one axis. For each experiment, uninjected and LacZ mRNA-injected embryos were used as negative controls.

### Luciferase reporter assay

100 pg β-catenin mRNA was co-injected with either 50 pg M50 Super 8X TOPflash or 50 pg M51 Super 8X FOPflash [35] into the animal pole region of *Xenopus* embryos at the 1-cell stage, and embryos were incubated at 20°C until stage 11-12 (middle gastrulae). Triplicate embryos were collected and washed in 0.1x Steinberg solution and homogenized in the luciferase assay system (E1500, Promega) following the manufacturer’s guidelines. Lysates were then transferred to individual wells in a 96-well plate, and luciferase activity was measured using a Centro LB960 microplate reader (Berthold Technologies). Statistical analysis was carried out with GraphPad Prism 9 software (San Diego, California USA, www.graphpad.com), and two-way ANOVA was used to analyze statistical significance (*P=0.05*).

### Microinjection for IP-MS analysis

β-catenin mRNA was injected into fertilized *X. laevis* embryos at the 1-cell stage. Due to differences in translation efficiency, the amount of mRNA injected for each metazoan FLAG-tagged β-catenin (3 ng of *B. mikado* versus 1 ng of *X. laevis*, *E. fluviatilis,* and *N. vectensis*) was optimized such that FLAG-tagged protein expression was roughly equivalent. Uninjected embryos were used as controls. Embryos were then cultured in 1x Steinberg Solution containing 5% Ficoll at 20°C until stage 12.

### Immunoprecipitation and western blot analysis

Homogenization and immunoprecipitation were as described [36] with slight modifications. Gastrula embryos (18 hours post-fertilization (hpf)) were lysed by pipetting in 1 mL ice-cold lysis buffer (20 mM Tris-HCl, pH 8, 70 mM KCl, 1 mM EDTA, 1% NP-40, 10% glycerol and 1× Roche complete proteinase inhibitor cocktail. This was followed by sonication in a water bath at 4°C for 60 sec. The lysate was then clarified by centrifugation for 15 min at 14000 rpm at 4°C to remove cell debris, pigment, and yolk. Supernatant was recovered, and subsequently, protein concentration was measured using a Direct Detect infrared spectrometer (Merck, Darmstadt, Germany). Aliquots of supernatant were then snap-frozen in liquid nitrogen and stored at –80°C. For immunoprecipitation, lysate containing 3 mg total protein was mixed with 40 μL anti-FLAG M2 magnetic beads (Sigma Aldrich, USA) in 1000 μL, and incubated overnight at 4°C. Beads were then washed three times with 500 μL cold lysis buffer and resuspended in 65 μL 2× sample buffer (Nacalai, Japan).

For western blot analysis, sample buffer was added to protein lysates or IP elutes and boiled at 95°C for 5 min. Subsequently, equivalent amounts of protein were separated on SDS-PAGE and transferred to a PVDF membrane (Biorad). The following antibodies were used for protein blotting: TUBA1A (Sigma-Aldrich, T6074, 1/1000), CTNNA1 (Santacruz, sc-9988, 1/500), FLAG-tag sequence (Sigma-Aldrich, F1804, 1/1000), and GAPDH (Santacruz, sc-47724, 1/1000). Signals were developed with horseradish peroxidase-conjugated goat against mouse secondary antibody (Jackson Immuno Research, 115035003, 1/10000) and chemiluminescence was developed by treating the membrane with ImmunoStar Zeta (Wako Pure Chemical Industries).

### In-gel trypsin digestion

45 μL of elute were loaded into lanes of a 7.5% polyacrylamide gel for SDS-PAGE. Protein bands were visualized using SimplyBlue SafeStain (Thermo Fisher Scientific). Using a clean scalpel, individual gel lanes were excised into six fractions from high to low molecular weight. These fractions were further diced into small pieces (2 mm^3^) and transferred to a low-binding, 96-well Eppendorf plate. Briefly, gel pieces were washed with MilliQ water, followed by 50 mM ammonium bicarbonate in 50% (vol/vol) acetonitrile and shrunken by adding 100% acetonitrile for 5 min. Proteins were reduced at 56°C with 160 μL of 10 mM dithiothreitol in 50 mM ammonium bicarbonate for 30 min and then alkylated with 160 μL of 55 mM iodoacetamide in 50 mM ammonium bicarbonate for 30 min in darkness. Finally, gel pieces were washed with 50 mM ammonium bicarbonate in 50% (vol/vol) acetonitrile, followed by 100% acetonitrile to complete dryness.

Gel pieces were rehydrated with 15 μL of 10 ng/μL trypsin in digestion buffer (50 mM ammonium bicarbonate containing 10% (vol/vol) acetonitrile) for 15 min. The liquid volume was adjusted with 60 μL of digestion buffer to prevent sample drying, and pieces were incubated overnight at 37°C. Digestion was stopped by adding formic acid to achieve a final concentration of 5%, and supernatant was transferred into a new 96-well plate. Tryptic peptides were extracted from the gel by adding 100 μL of 50% acetonitrile/ 5% formic acid (vol/vol) for 45 min, followed by 100 μL of 100% acetonitrile for 5 min. Extracted peptides were pooled in the 96-well plate and concentrated by vacuum centrifugation at 42°C for 2 h using the HPLC method of a Genevac EZ-2 Elite (ATS) vacuum evaporator. Dried peptides were desalted using C_18_ StageTips [75]. Briefly, C_18_ StageTips were prepared by packing C_18_ solid phase extraction disks (CDS Empore^TM^) in 200 μL tips, activated with 100 μL of 80% acetonitrile/ 0.1% formic acid (vol/vol) and conditioned with 200 μL of 1% acetic acid/ 0.5% formic acid (vol/vol). Peptides were resuspended in 50 μL of 1% acetic acid/ 0.5% formic acid (vol/vol) and then loaded onto C_18_ StageTips. After washing with 200 μL of 0.1% formic acid, peptides were eluted with 100 μL of 80% acetonitrile/ 0.1% formic acid and dried using the vacuum evaporator as described above.

### LC-MS/MS analysis

Dried peptides were resuspended in 25 μL of 5% methanol/ 0.1% formic acid in MilliQ water (vol/vol), and separated with an ultrahigh-performance liquid chromatography system (Waters nanoACQUITY UPLC, Waters) on a trap column (2 cm x 180 μm, nanoEase M/Z Symmetry C_18_ Trap Column, Waters), and a separation column (15 cm x 75 μm, nanoEase M/Z HSS T3 Column, Waters). Solvent A was 0.1% formic acid and solvent B was 0.1% formic acid in 80% acetonitrile. The column temperature was set to 40°C. The UPLC system was coupled online to an Orbitrap Fusion Lumos mass spectrometer (Thermo Fisher Scientific). For each sample, 5 μL were injected, and peptides were separated at a flow rate of 0.5 μL/min using a one-hour gradient of 1-50% solvent B. The mass spectrometer was set to perform data acquisition in positive-ion mode at a resolution of 120K, MS1 scan range between 400 and 1500 m/z, a maximum ion accumulation time of 50 msec with an AGC target of 4.0e5. Other settings included 3 sec between master scans (MS1), activation of monoisotopic peak determination of peptide (MIPS), and an ion isolation window of 1.2 m/z. MS2 spectra were analyzed either by the Orbitrap or the linear ion trap using the CHOPIN method [76], depending on the charge and intensity of the peak. Precursor ions carrying 2+ charges were fragmented with collision-induced dissociation (CID) in the ion trap at a collision energy of 35%. Precursor charges from 3+ to 7+ with precursor intensity greater than 5.0^e5^ were fragmented with High Collision Dissociation (HCD) in the orbitrap at a collision energy of 25% and the orbitrap resolution set at 15K. The remaining ions, with charges from 3+ to 7+ with precursor intensity less than 5.0^e5^, were fragmented with CID as above. Dynamic exclusion was applied after 1 event for 12 sec with a mass tolerance of 10 ppm. All raw LC-MS/MS data have been deposited at the ProteomeXchange Consortium (http://proteomecentral.proteomexchange.org) via jPOST with the data set identifier <PXD053223>. Proteome Discoverer output is also available on ProteomeXchange Consortium with the same identifier as above.

## Data analysis

All MS and MS/MS spectra were analyzed with Proteome Discoverer software V2.2 (Thermo Fisher Scientific, Inc) with the SEQUEST HT search engine for protein identification and label-free quantitation. Database searches were performed against the Uniprot *X. laevis* database (11/Dec/2019; 57070 entries) and the common Repository of Adventitious Proteins (cRAP; http://www.thegpm.org/crap/). Search parameters for identification were trypsin enzyme, allowing up to 2 missed cleavages, with precursor and fragment mass tolerance set to 20 ppm and 0.8 Da, respectively. Carboxyamidomethylation of cysteine was set as a fixed modification, while methionine oxidation, asparagine, glutamine deamidation, and N-terminal acetylation were variable modifications. Results were filtered using a False Discovery Rate (FDR) of <1% as a cutoff threshold at the protein level, determined by the Percolator algorithm in Proteome Discoverer software. Protein abundances in each fraction were summed and normalized using the “Total Peptide Amount” setting. For IP samples, protein ratio calculations were performed using pairwise ratios and proteins identified in at least two of three replicates were included in the statistical analysis. Differences in protein composition was evaluated using background-based ANOVA analysis, as implemented in Proteome Discoverer. Proteins were considered significantly changed if the log2 fold change in abundance between the control (uninjected) and sample was greater than 1, with an adjusted p value less than 0.05.

## Availability of data and materials

The dataset supporting conclusions of this article is available in the ProteomeXchange Consortium (http://proteomecentral.proteomexchange.org) via the jPOST partner repository (https://jpostdb.org) [92] under accession number JPST001441. Other datasets and plasmids used in this study are available from the corresponding author upon request.

## Declarations

### Ethics approval

All experiments and procedures carried out using *Xenopus laevis* were approved by the Okinawa Institute of Science and Technology Graduate University’s Animal Care and Use Committee (Approval ID: 2020-308-2).

### Consent for publication

Not applicable.

## Acknowledgements

We thank the OIST Instrument Analysis Section for assistance with mass spectrometry experiments and the Animal Resource Section for the assistance in rearing *Xenopus* frogs. We thank Dr. Steven D. Aird (https://www.sda-technical-editor.org/) for editing the manuscript. This work was supported by OIST Graduate University and Life Science Research (BINDS)) from AMED under Grant Number JP21am0101110 (support number 0883). This work was supported by JSPS KAKENHI Grant Number JP19K06796 (EH), JP22K06348 (YY), and JP20K06662 (HW).

## Author contributions

IM, HW conceived and designed this study. IM carried out most experiments and analyzed data. CK, AVB, YH, HE carried out proteomic experiments and data analysis. YT, KT performed 3D modelling. YY instructed IM on experiments using *Xenopus* embryos. IM, CK, KT, HW wrote the manuscript. All authors have read and approved the manuscript.

## Figure Legends

**Figure S1.**
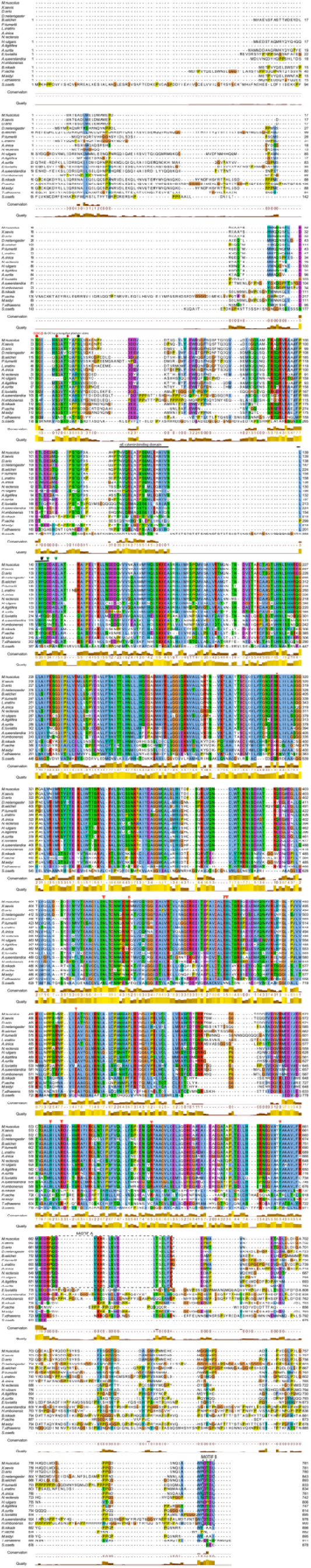
Alignment of metazoan β-catenin and *S. rosetta* ARM6 proteins. Red and black circles represent GSK3β and CK1α phosphorylation sites, respectively. The black, bold line represents the α-catenin binding motif. The black arrow represents a tyrosine residue critical for binding α-catenin and green arrows represent acidic residues that promote or inhibit this interaction. Red asterisks represent lysine residues vital to binding of both TCF and E-cadherin. Red arrows represent residues key in binding TCF. Motifs A and B in the C-transactivation domain are labelled with dotted boxes.

**Figure S2.**
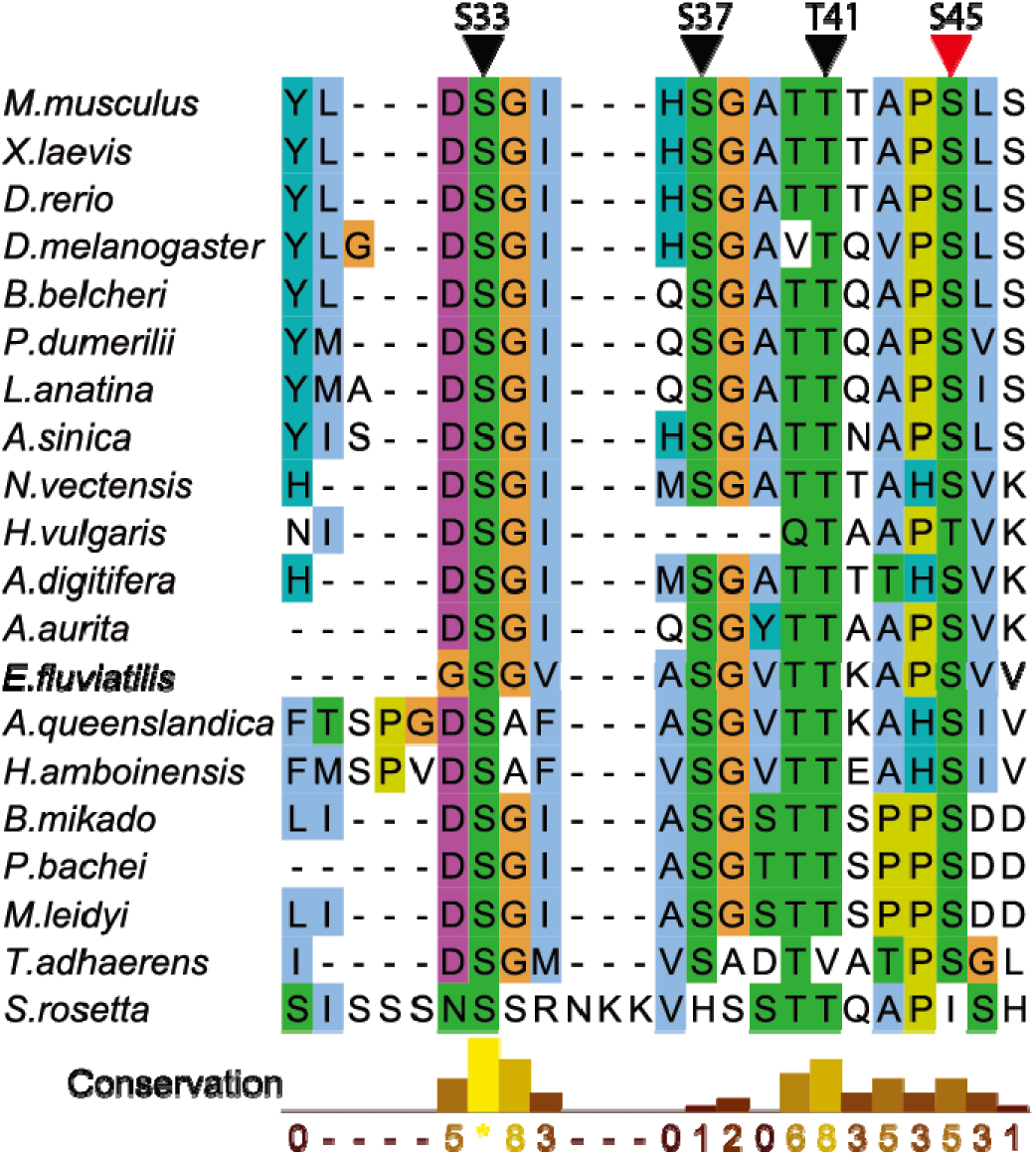
Assumed GSK3β and CK1α phosphorylation sites of nonbilaterian β-catenins. All three GSK3β phosphorylation sites (black arrowheads) were conserved in several metazoans, except for *H. vulgaris* and *T. adhaerens*. Interestingly, the choanoflagellate, *S. rosetta* ARM6 showed conservation of two sites corresponding to S37 and T41. The CK1α phosphorylation site (red arrowheads) was conserved among metazoans, but not in *S. rosetta* ARM6.

**Figure S3.**
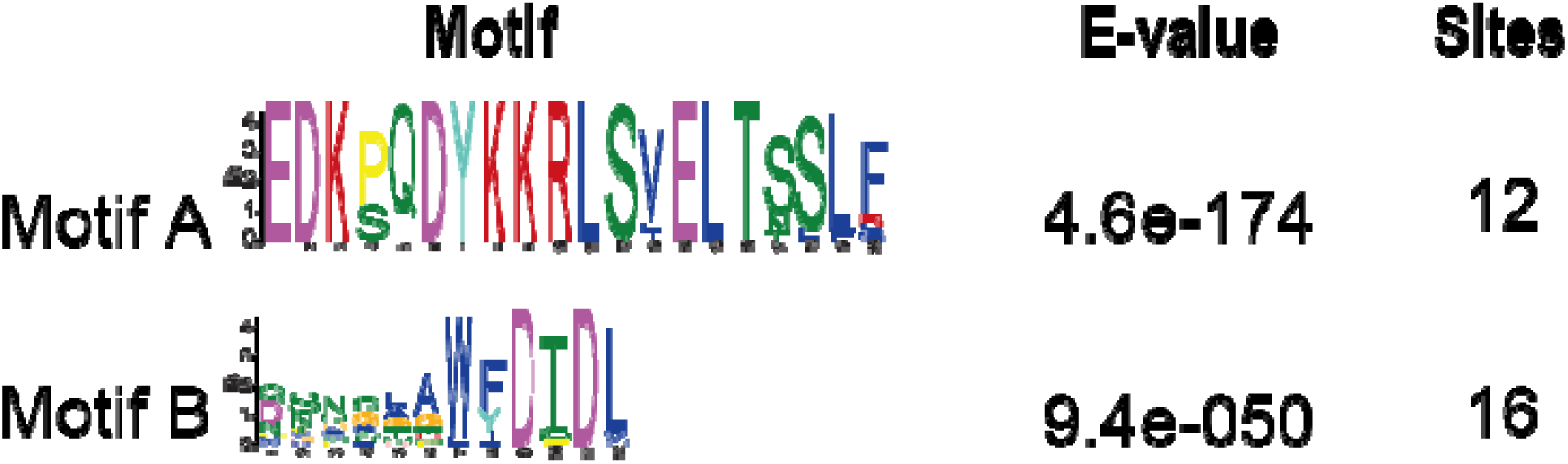
Multiple Expectation maximization for Motif Elicitation (M 815 EME) analysis. Two statistically significant motifs were identified in the C-terminal region. The sequence of each motif is shown in larger letters reflecting its significance.

**Figure S4.**
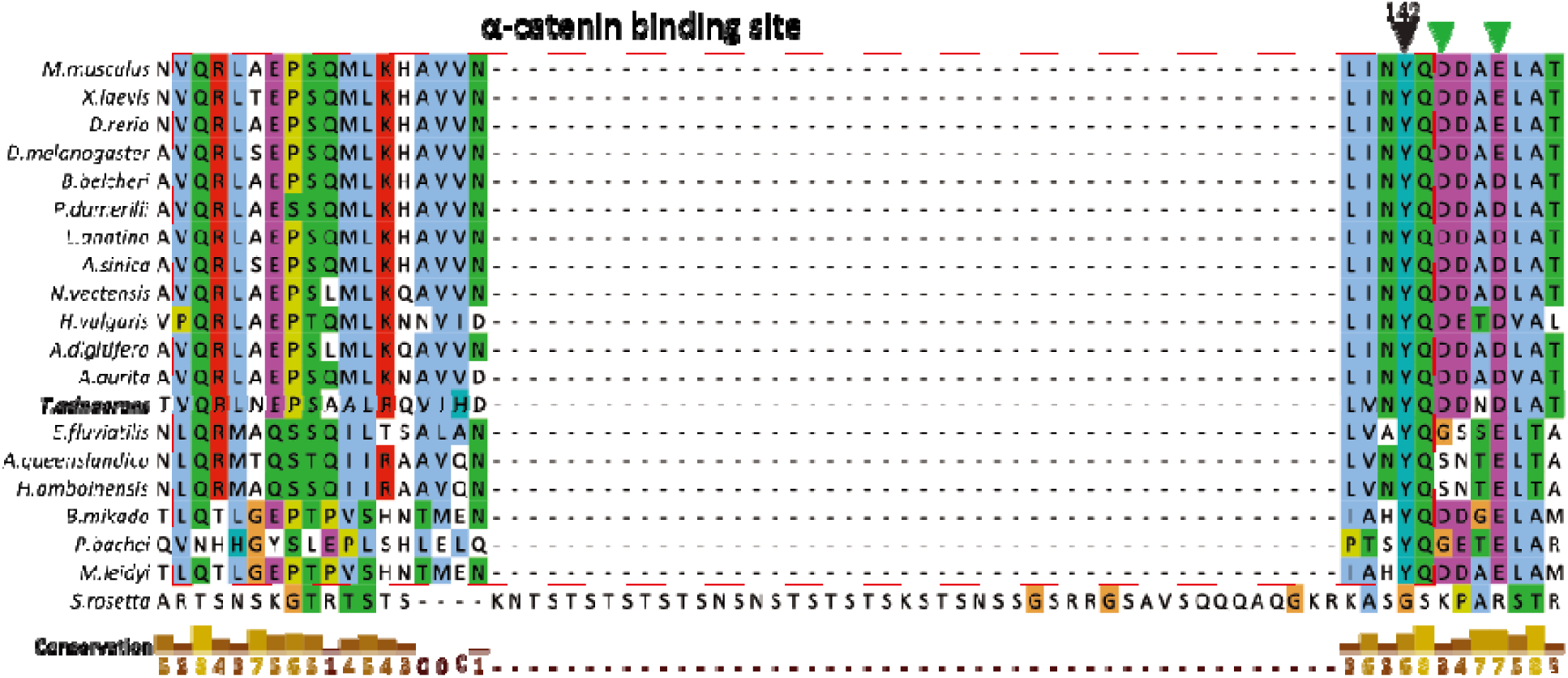
Sequence alignment of α-catenin binding sites of β-catenins. Although conservation across the entire α-catenin binding region varied, the critical tyrosine residue (black arrowhead) was conserved in metazoans. The two acidic residues (green arrowheads) that affect binding of α-catenin are also conserved in metazoans, with the exception of poriferans.

**Figure S5.**
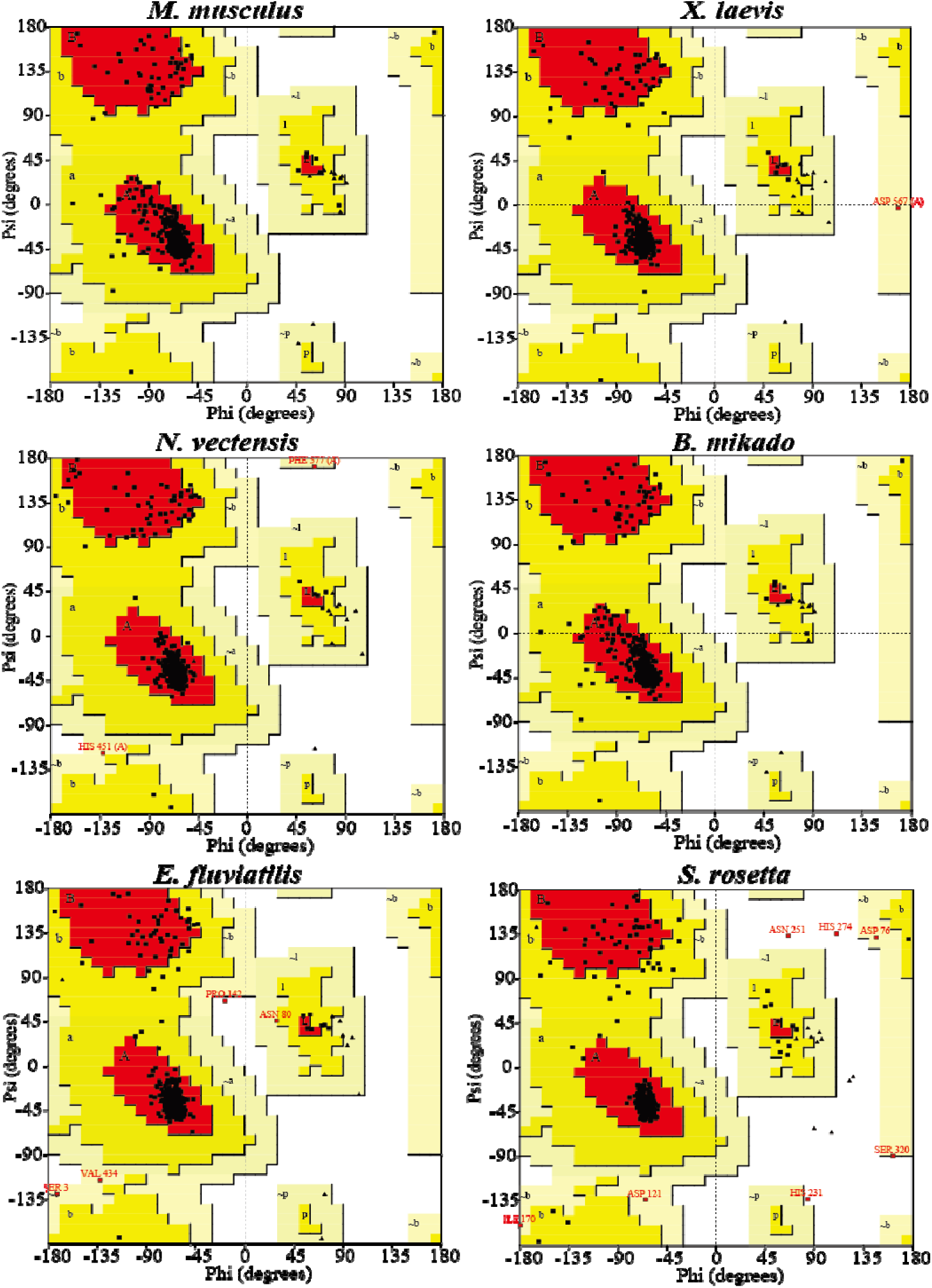
Ramachandran plot analysis. The majority of residues (black squares) were localized in favored positions (red regions), followed by those in allowed positions (yellow regions). Few residues were localized in generously allowed positions (light yellow). Disallowed regions are indicated in white.

**Figure S6.**
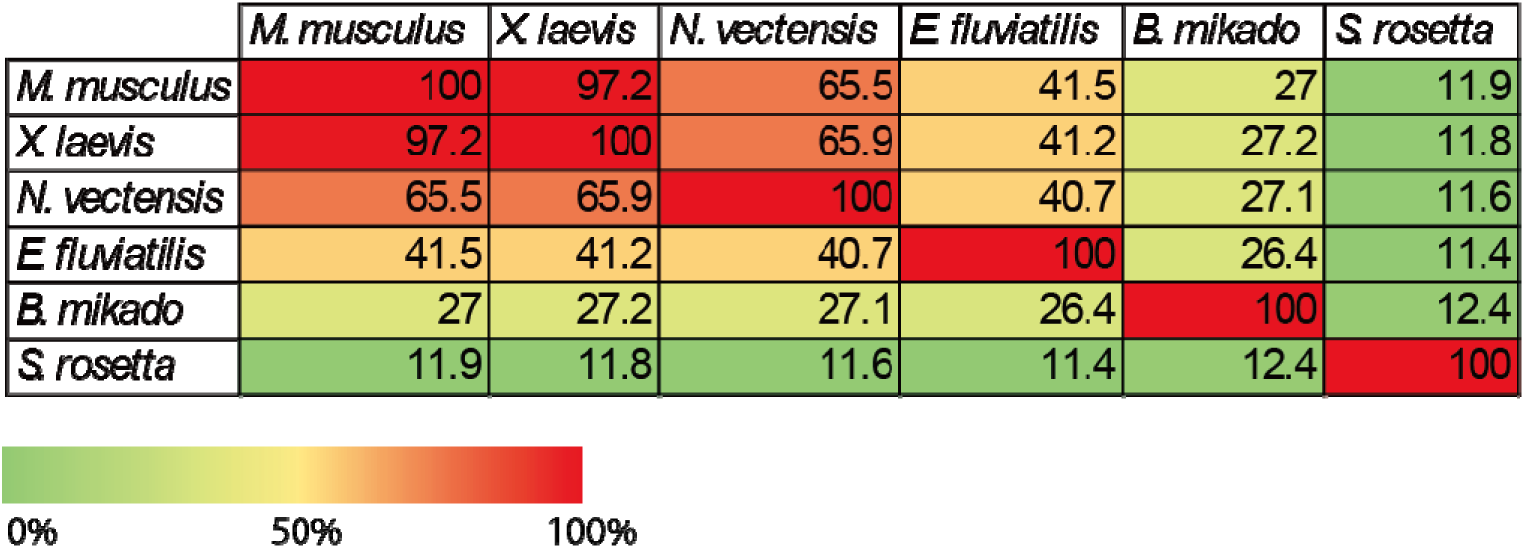
Percentage amino acid sequence similarity between metazoan β-catenins. Sequence similarity was high in Bilateria and Cnidaria (>50%), but low in Porifera (41%) and Ctenophora (27%). Non-metazoan *S. rosetta* ARM6 protein displayed the lowest similarity (12%) to mouse β-catenin.

**Figure S7.**
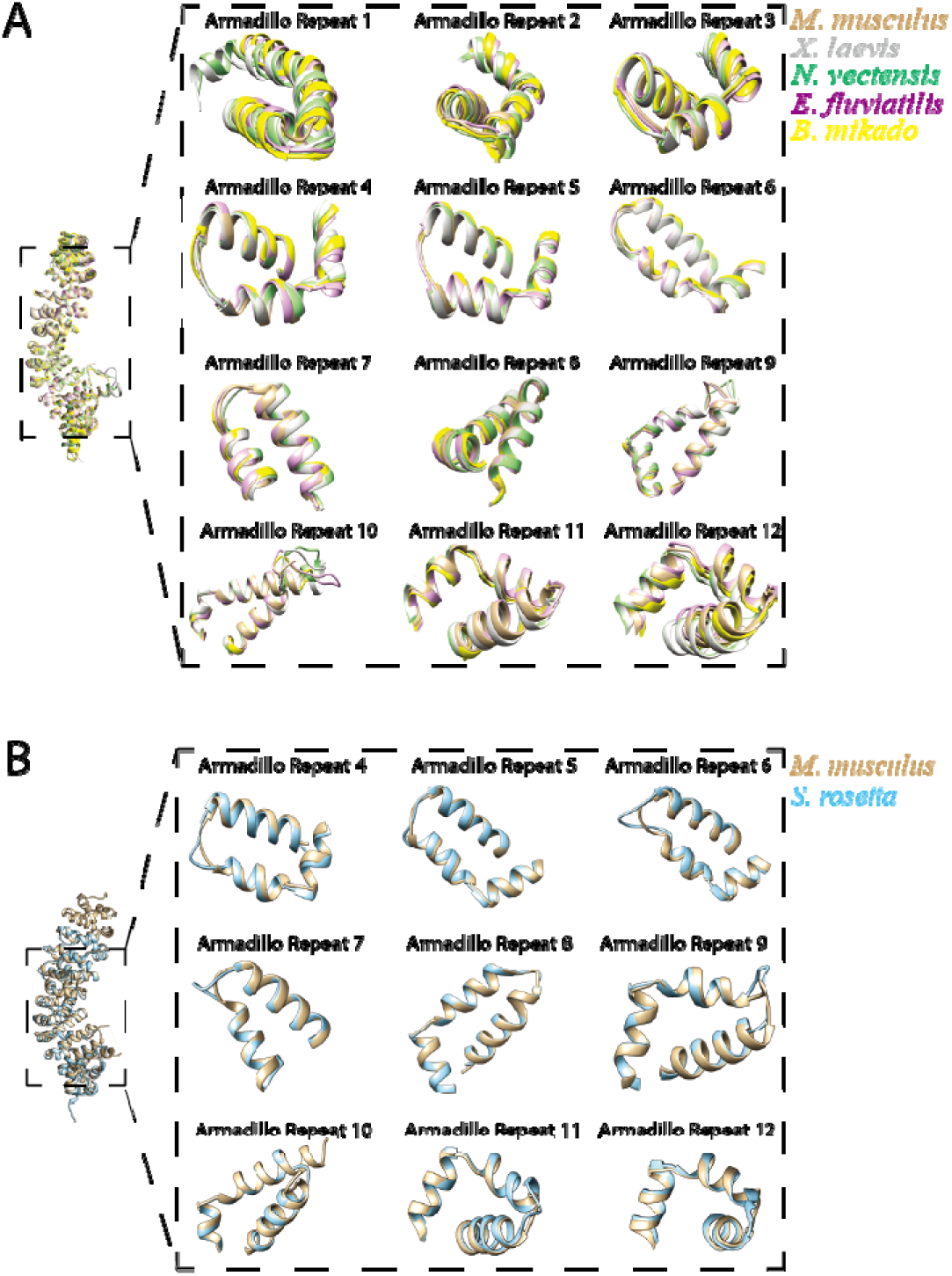
Conservation of the central region of metazoan β-catenins. Superimposition of predicted β-catenin structures onto template *M. musculus* β-catenin showed that model metazoan β-catenin had twelve conserved armadillo repeats while (A) while *S. rosetta* ARM6 protein had only nine (B).

**Figure S8.**
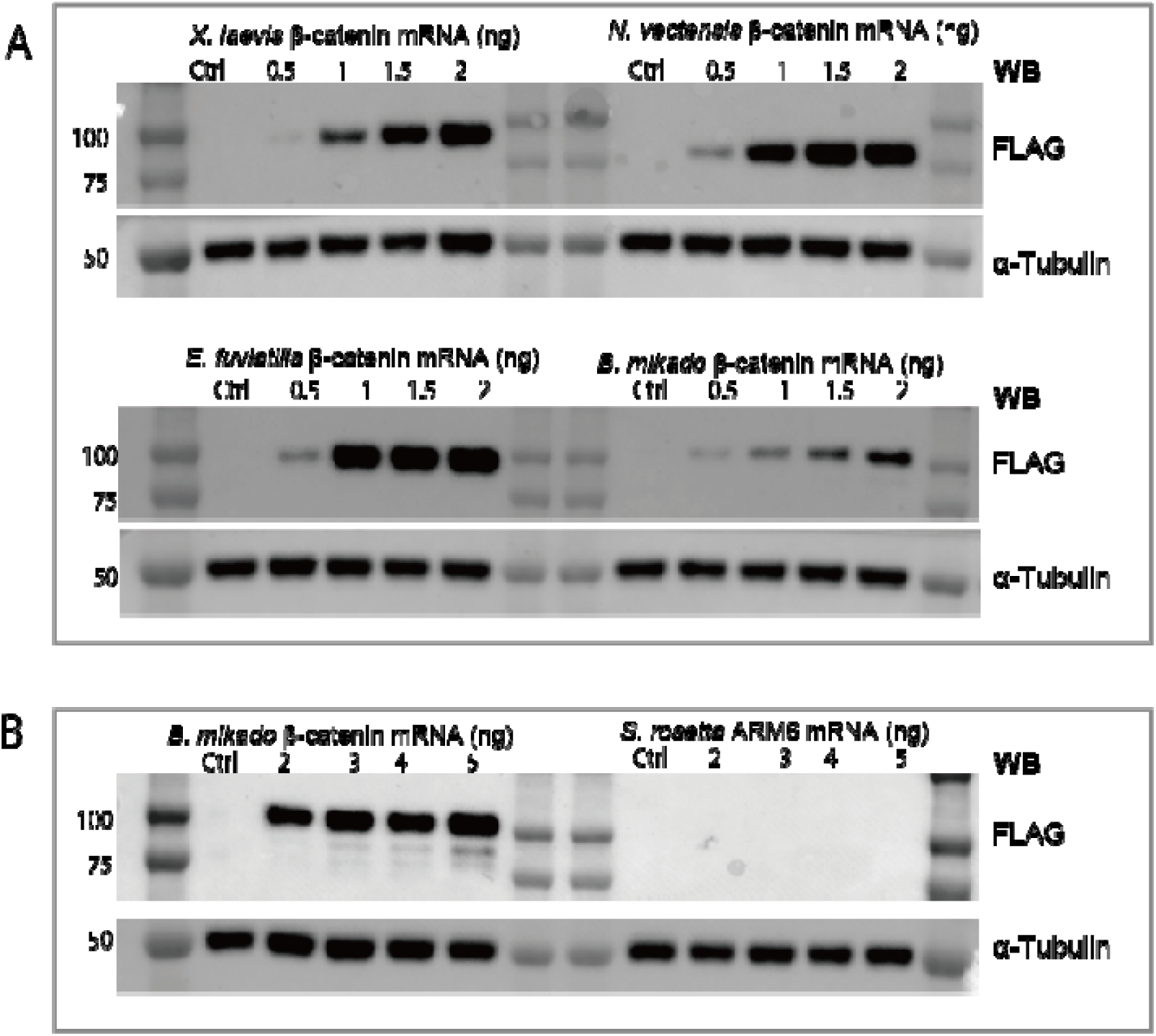
Expression of exogenous β-catenin protein in *Xenopus* embryos. (A) Western blot of *Xenopus* and nonbilaterian FLAG-tagged β-catenin proteins after injection of increasing concentrations of mRNA into *Xenopus* eggs. (B) No expression of *S. rosetta* ARM6 was observed, even in embryos injected with a high concentration of the mRNA.

**Figure S9.**
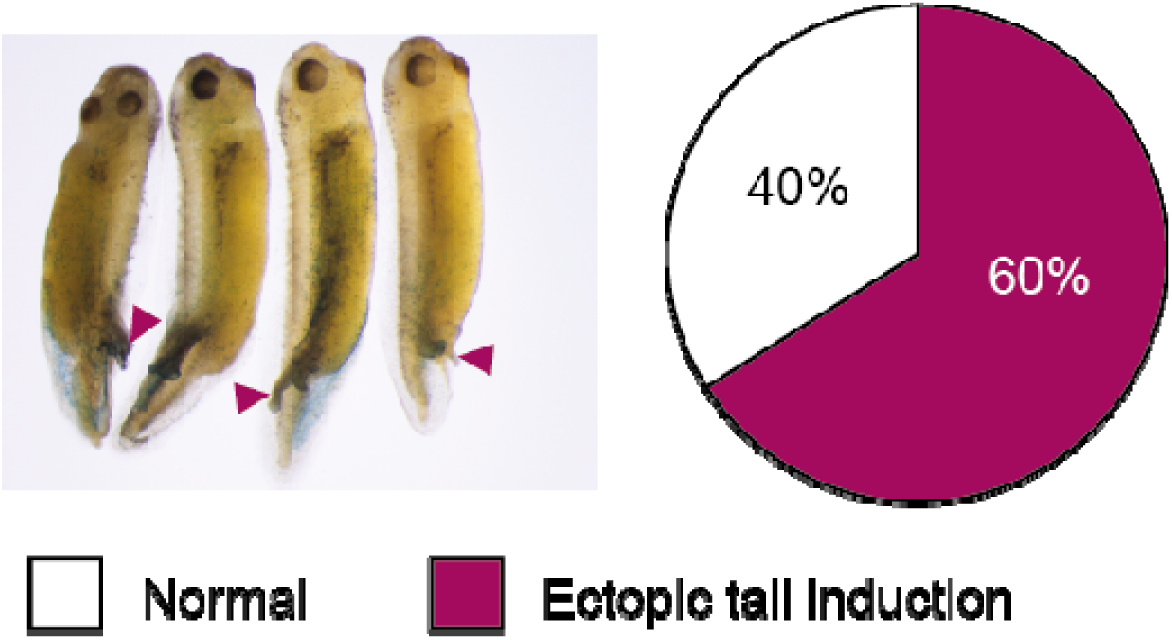
Unique structure provoked by ectopic expression of ctenophore β-catenin in Xenopus embryos. Injection of B. mikado β-catenin in X. laevis embryos (n=20) resulted in an ectopic tail-like structure (red arrowheads) at high concentrations of mRNA (500 pg).

**Figure S10.**
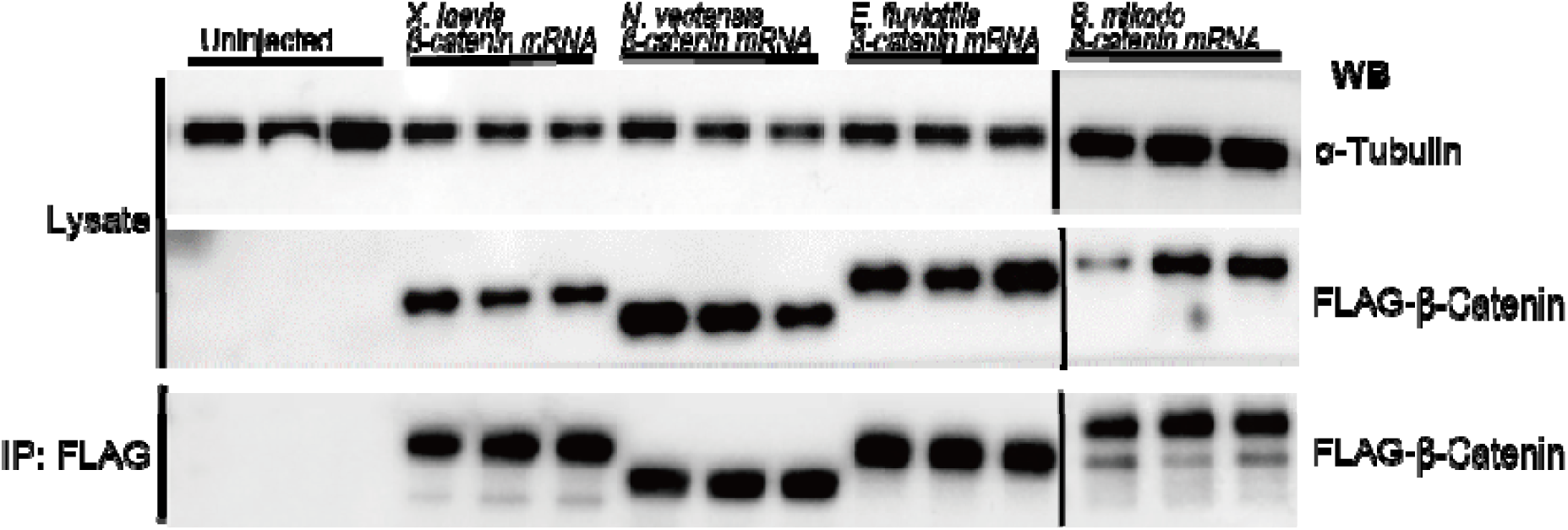
Expression and immunoprecipitation of FLAG-tagged β-catenin proteins. Western blotting showing expression levels of α-tubulin (internal standard) (top) and β-catenin (middle) in embryos injected with β-catenin mRNA and immunoprecipitated FLAG-β-catenin (bottom). Three replicates are shown for each β-catenin construct.

**Figure S11.**
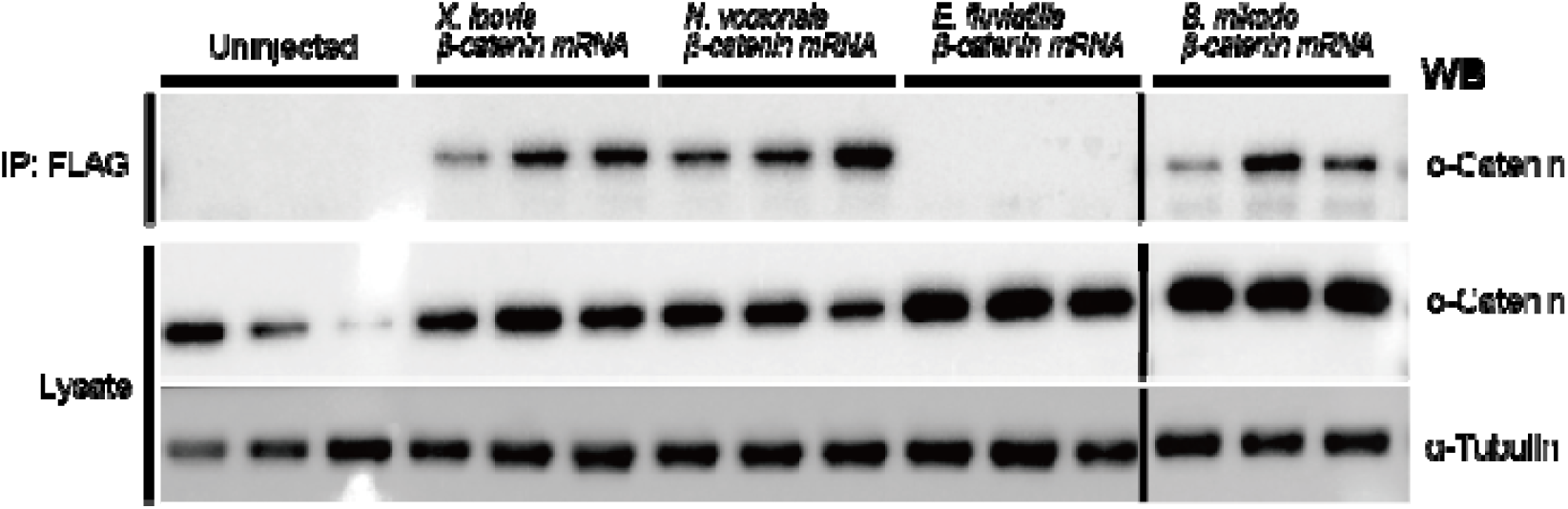
Co-immunoprecipitation of *Xenopus* α-catenin with FLAG-tagged β-catenin proteins. Western blotting showing expression levels of α-catenin (middle) and α-tubulin (bottom) (internal standard) in embryos injected with β-catenin mRNA and α-catenin co-immunoprecipitated with FLAG-β-catenin (top). Three replicates are shown for each β-catenin construct.

**Figure S12.**
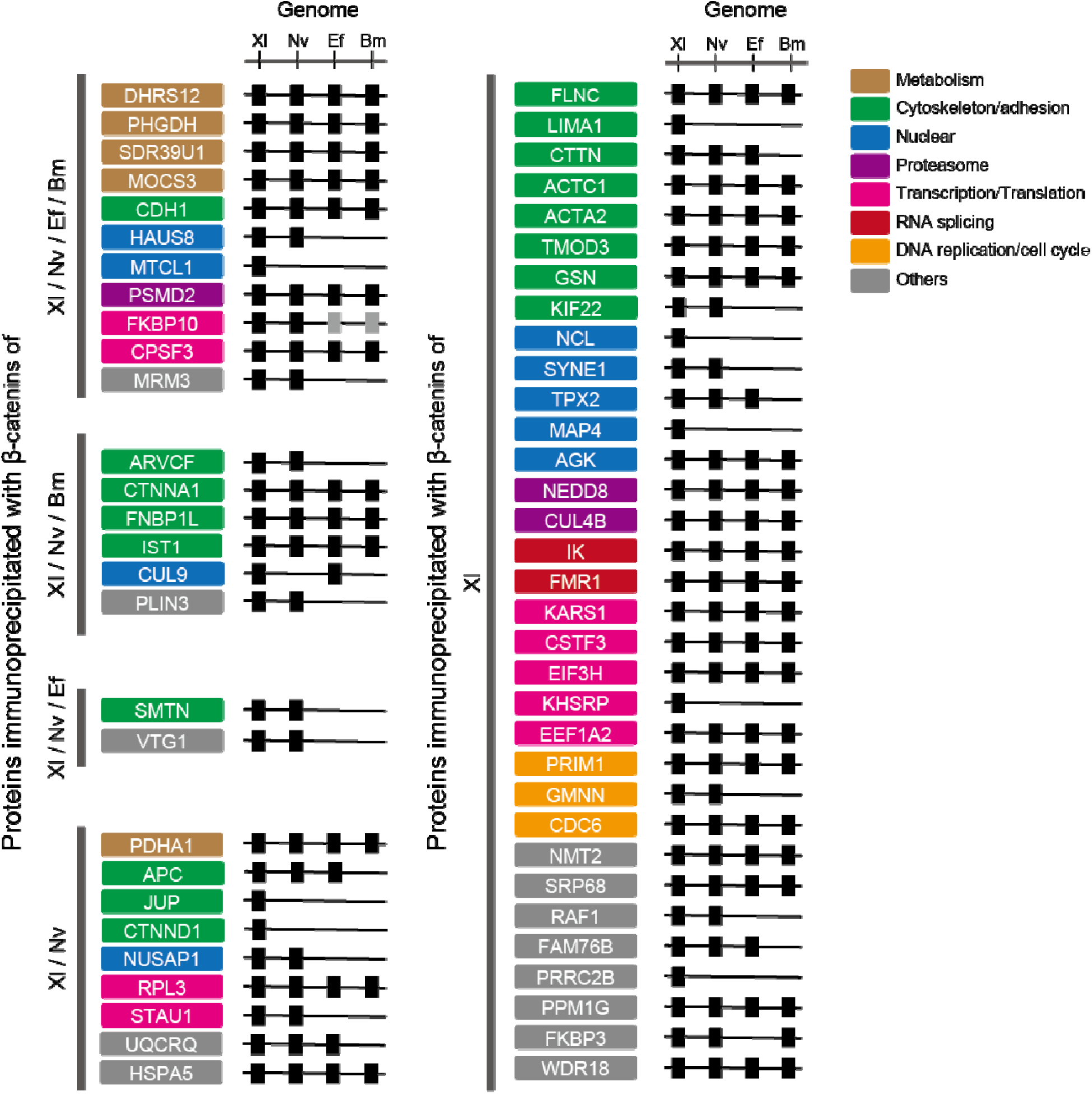
Distribution of β-catenin complex protein homologs in bilaterian and nonbilaterian genomes. Black and shaded boxes indicate homologs with high and low confidence, respectively.

**Table S1.** Xenopus proteins co-immunoprecipitated with metazoan β-catenins.

**Table S2.**
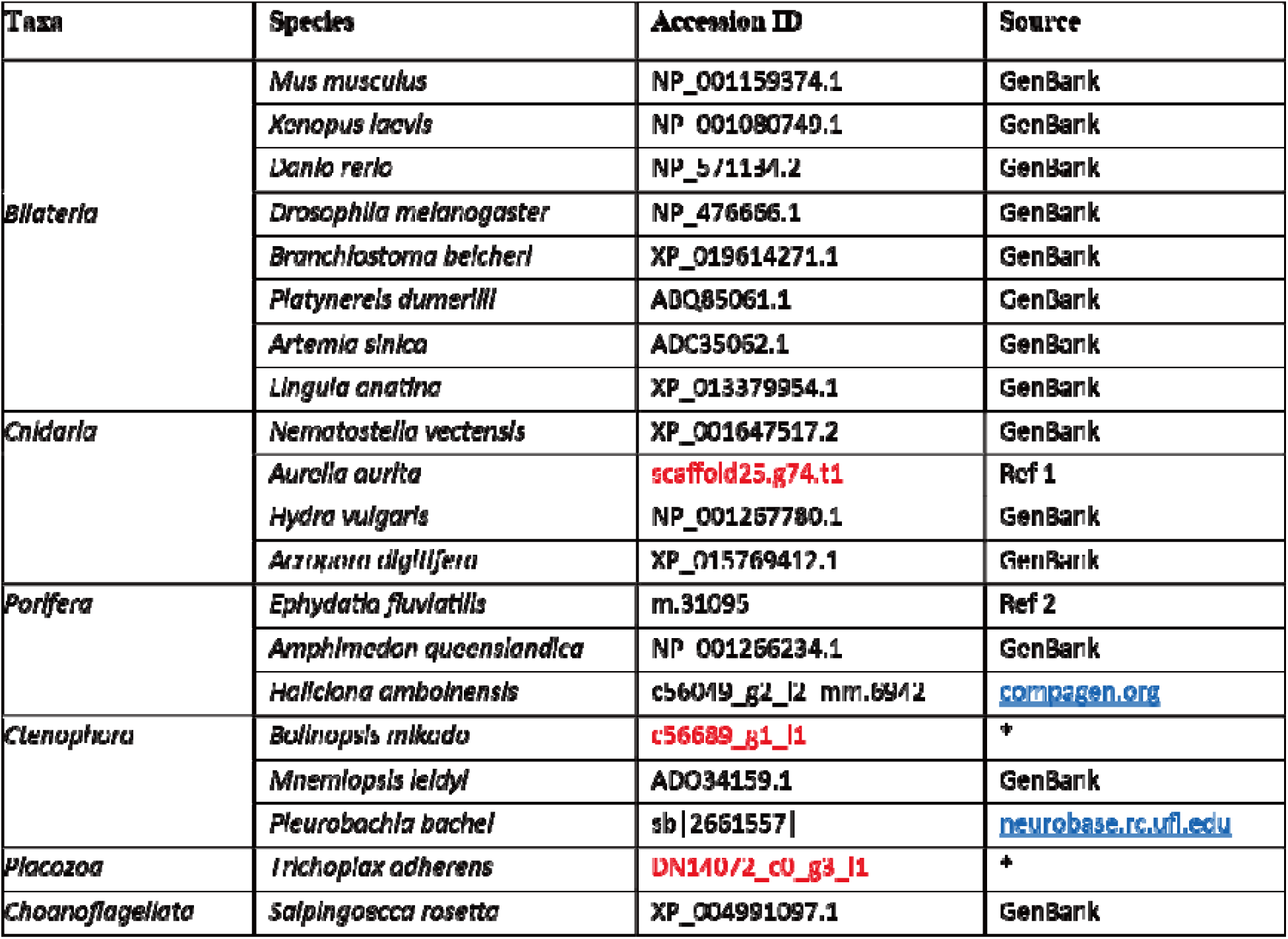
Accession IDs in red were translated to respective protein sequences using ExPASy (web.expasy.org/translate/). Asterisks, transcriptomic analysis carried out by our group.

**Table S3.**
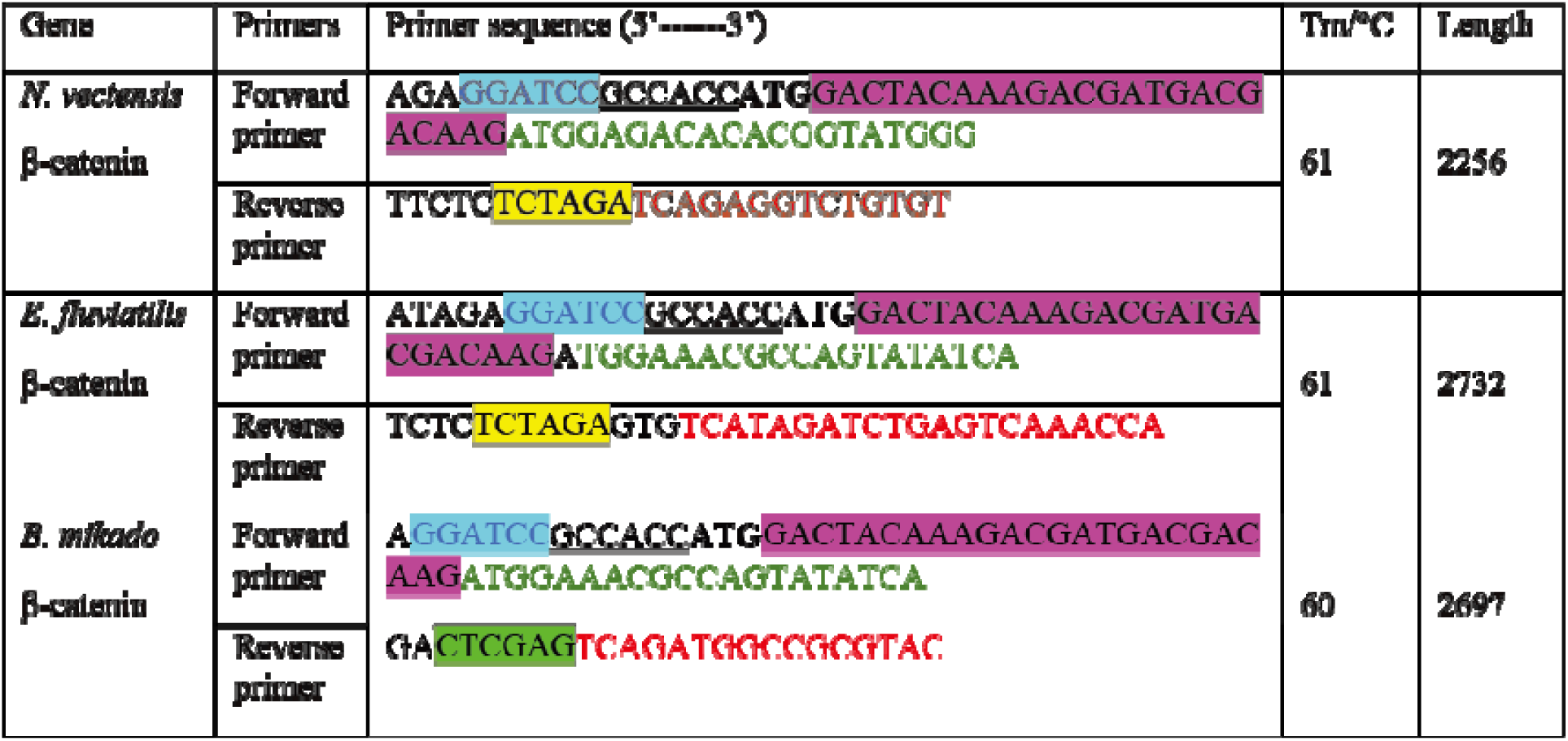
Primers used in cloning β-catenin. Restriction digest sites (highlighted blue (BamH1), yellow (Xba1) and green (Xho1). A flag-tag coding sequence (highlighted pink) was also included in the forward primer. Green and red nucleotides represent binding sites for forward and reverse primers, respectively. Bold “ATG” represents the start codon. A Kozak sequence (underlined) was also included before the start codon to improve translation initiation. *X. laevis* β-catenin and *S. rosetta* ARM6 were generated by FASMAC, Japan.

Table S4.

| Plasmid | Description | Source |
| --- | --- | --- |
| pCS2+ Flag-Nvec-β-cat | Flag-tagged *N. vectensis* β-catenin | This study |
| pCS2+ Flag-Xlae-β-cat | Flag-tagged *X. laevis* β-catenin | This study |
| pCS2+ Flag-Eflu-β-cat | Flag-tagged *E. fluviatilis* β-catenin | This study |
| pCS2+ Flag-Bmik-β-cat | Flag-tagged *B. mikado* β-catenin | This study |
| pCS2+β-gal | β- galactosidase | Yuuri Yasuoka |
| *pCSf107_8x Super TOPFLASH | Beta-catenin reporter. TCF/LEF sites upstream of a luciferase reporter. | Veeman et al. (2003) |
| *pCSf107_8x Super FOPFLASH | Contains mutated TCF binding sites upstream of a luciferase reporter. | Veeman et al. (2003) |

\*pCSf107 is vector that contains SP6/T7 terminator sequences (Mii and Taira, 2009).

